# Apical Size Reduction by Macropinocytosis Alleviates Tissue Crowding

**DOI:** 10.1101/2024.08.05.606662

**Authors:** Enzo Bresteau, Eve E Suva, Christopher Revell, Osama A Hassan, Aline Grata, Jennifer Sheridan, Jennifer Mitchell, Constandina Arvantis, Farida Korobova, Sarah Woolner, Oliver Jensen, Brian Mitchell

## Abstract

Tissue crowding represents a critical challenge to epithelial tissues, which often respond via the irreversible process of live cell extrusion. We report cell size reduction via macropinocytosis as an alternative mechanism. Macropinocytosis is triggered by tissue crowding via mechanosensory signaling, leading to substantial internalization of apical membrane and driving a reduction in apical cell size that remodels the epithelium to alleviate crowding. We report that this mechanism regulates the long-term organization of developing epithelium in response to proliferation-induced crowding but also serves as an immediate response to acute external compression. In both cases, inhibiting macropinocytosis induces a dramatic increase in cell extrusion suggesting cooperation between cell extrusion and macropinocytosis in response to compression. Our findings implicate macropinocytosis as an important regulator of dynamic epithelial remodeling.

## INTRODUCTION

Epithelial tissue is a conserved feature of all metazoan animals, and its barrier function is essential for multicellular life. It can serve as a mechanical barrier between organs that is capable of withstanding a variety of stresses, including physical compression from the environment as well as the significant tissue remodeling that occurs during morphogenesis. Epithelial tissue crowding, the accumulation of epithelial cells within a confined space, typically arises from cell proliferation but can be exacerbated by externally applied compression. Defects in how tissues respond to crowding are associated with aberrant cell division and tumor growth ^1-4^. To alleviate tissue crowding, an epithelium must remove some of its apical surface. An important and well-studied response is cell removal via live cell extrusion ^1, 2, 4^. Here we propose that epithelial crowding can be addressed without irreversible cell loss, by multiple cells reducing their apical size. We propose that membrane internalization via macropinocytosis drives apical size reduction in to alleviates tissue crowding.

Macropinocytosis is a non-specific endocytic pathway that relies on the formation of large circular actin ruffles that protrude from the cell surface, leading to the engulfment of extracellular material into large vesicles ^5, 6^. Macropinocytosis has typically been associated with nutrient acquisition, receptor recycling, and immune surveillance ^5^; however, recent studies have also implicated macropinocytosis in membrane remodeling ^7, 8^. More broadly, membrane reshaping by various forms of endocytosis has been shown to influence stress response and morphogenesis ^9-12^. However, the role of endocytic processes in regulating tissue mechanics remains largely unknown.

In this study, we utilize the surface epithelium of developing *Xenopus* embryos to propose a previously unappreciated role for membrane internalization via macropinocytosis as a mechanism for apical size reduction in response to tissue crowding. First, we report that macropinocytotic events increasingly occur during the growth of the epithelium to alleviate local crowding during development. Additionally, we propose that macropinocytosis can serve as an immediate response to environmental stress. We report that waves of macropinocytotic events occur to remodel the epithelium in response to externally induced compression of the epithelium. Altogether, our results suggest that macropinocytosis cooperates with live cell extrusion in response to tissue crowding during both environmental stress and epithelial morphogenesis.

## RESULTS

### Constitutive macropinocytosis occurs in the apical surface of the Xenopus embryonic epidermis

Actin ruffling has been reported to occur in a wide range of tissues under diverse physiological conditions. Actin staining as well as imaging of dynamic actin using Lifeact in late Stage (ST34+) *Xenopus* embryos revealed numerous events of circular actin ruffling on the epithelial surface **(Fig. 1A, 1B, Movie S1, S2)**. These events initiate rapidly and terminate in 9.3 (±2.3) min **(Fig. 1D)**. The actin ruffles are quite large, with a maximal area of 94 (±36) μm^2^ **(Fig. 1E)**, which represents 22.8% (±6.6%) of the total apical surface of the host cell **(Fig. 1F)**. While these ruffles can occur anywhere on the apical surface of the cell, we find that they preferentially occur close to cell-cell junctions **(Fig. 1G)**. Imaging embryos using scanning electron microscopy (SEM) revealed multiple phases of these events including circular ruffles protruding apically out of the cell as well as circular protrusions that appear to be closing in on themselves **(Fig. 1C)**. These structures are similar to the actin ruffles observed during macropinocytotic events that have been described in cell culture as well as other model organisms ^6, 13, 14^. To confirm that our observed actin ruffles lead to macropinocytosis we bathed embryos in media containing fluorescent dextran and found that the closing of circular actin ruffles indeed leads to the internalization of dextran-filled vesicles inside the cell **(Fig. 1H, Movie S3)**. Importantly, this occurs via the internalization of a substantial portion of the apical membrane to form a large vesicle inside the cell **(Fig. 1I)**.

**Fig. 1.**
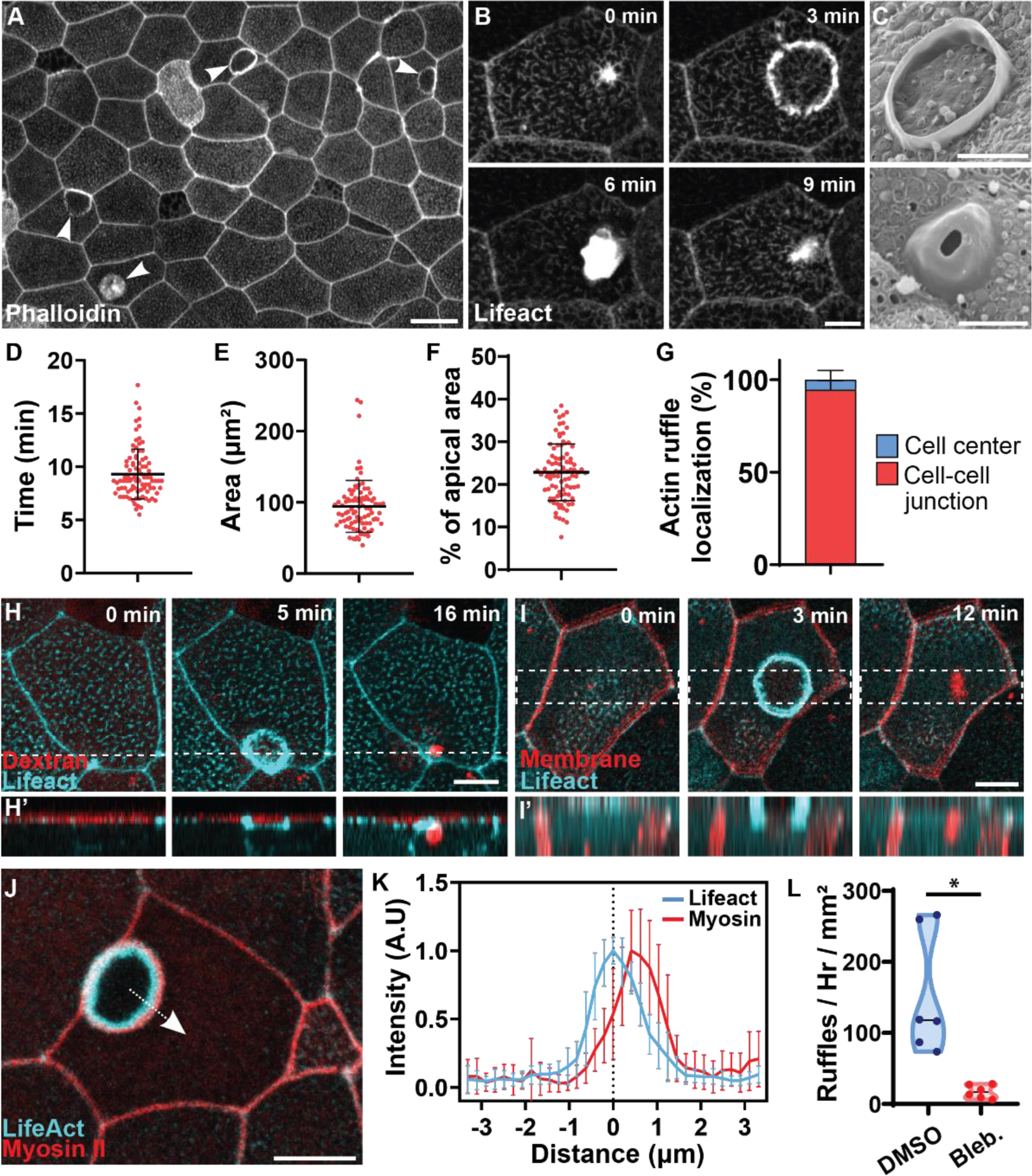
Constitutive macropinocytosis occurs in the apical surface of the *Xenopus* embryonic epidermis. (**A**) Actin ruffles (arrowheads) in the epidermis of *Xenopus* embryo stained with phalloidin. Scale bar 20 μm. (**B**) Timelapse of a single ruffle event in the epidermis of *Xenopus* embryo expressing Lifeact-fluorescent protein (FP). Scale bar 10 μm. (**C**) SEM images of actin ruffles as circular (top) or dome-shaped (bottom). Scale bar 5 μm. (**D**-**F**) Quantification of actin ruffle duration (D), peak area (E), and percentage of apical area (F) (n = 3 exp. and 90 ruffles). (**G**) Quantification of actin ruffle localization within the cell (n = 9 exp. and 361 ruffles). (**H**) Time lapse image of the Lifeact-FP (cyan) showing macropinocytotic internalization of fluorescent dextran (red) from the media. H’ is a z projection of the dotted line. Scale bar 10 μm. (**I**) Time lapse image of the Lifeact-FP (cyan) showing internalization of apical membrane (red) during macropinocytosis. I’ is a z projection of the dotted area. Scale bar 10 μm. (**J**) Circular actin ruffle (cyan, Lifeact-FP) surrounded by an outer ring of activated myosin II (red; SF9 intrabody). The dotted arrow shows an example of linescan averaged in (K). Scale bar 10 μm. (**K**) Localization of Lifeact and activated myosin II at the peak of an actin ruffles (n = 2 exp. and 15 ruffles). (**L**) Quantification of ruffle number in embryos treated with DMSO or Blebbistatin (n = 3 exp. and 6 embryos).

Myosins are often the driving force in generating dynamic actin-based structures ^15^. Imaging with a fluorescently tagged Myosin II intrabody (SF9) ^16^ revealed that active Myosin II localizes outside of the circular actin ruffles **(Fig. 1J, 1K, Movie S4)**, suggesting a role for actomyosin forces in the dynamic ruffle protrusion. Accordingly, treatment with the Myosin inhibitor, Blebbistatin, resulted in a complete loss of these macropinocytotic events **(Fig. 1L)**.

### Macropinocytosis is associated with tissue crowding and low levels of tension

We observe that macropinocytosis progressively increases during development. While we rarely observe macropinocytotic events in ST34 embryos, their occurrence is dramatically increased by ST42 **(Fig 2A, 2B)**. As development progresses, proliferation within the confined epithelium leads to an increase in cell density, which correlates with higher levels of macropinocytosis **(Fig 2A, 2B)**. To test the importance of cell density on macropinocytosis, we inhibited cell proliferation with the Cyclin Dependent Kinase (CDK) inhibitor Roscovitine ^17^. In control embryos we continue to see high levels of macropinocytosis at ST47. However, when we inhibit proliferation with Roscovitine, we see a significant decrease in macropinocytosis, suggesting that these events are induced in crowded tissues (**Fig 2C**). Tissue crowding has been associated with low levels of junctional tension in the epithelium ^2, 18-21^, suggesting that tension levels could modulate macropinocytosis. Additionally, membrane tension is a known regulator of endocytosis in general ^22^ and has recently been linked to macropinocytosis ^14, 23^. To address the importance of tensile forces on macropinocytosis, we utilized ectodermal tissue excised from ST10 embryos. This tissue can be cultured into a 3D epithelial organoid, or it can be plated onto fibronectin coated glass where it will adhere to the substrate and spread out into a 2D explant **(Fig 2D)**. The 3D organoids are generally considered to have lower tensile strength than explanted tissue due to the lack of an underlying substrate ^24-27^. As expected, low-tensile organoids have a higher cell density than the high-tensile explanted tissue **(Fig 2E, 2F, Movie S5, S6)**. We quantified macropinocytotic events and found that organoids have a significantly higher level of macropinocytosis as compared to the explants **(Fig 2E-F, Movie S5, S6)** consistent with roles for tensile force and tissue crowding in driving macropinocytosis.

**Fig. 2.**
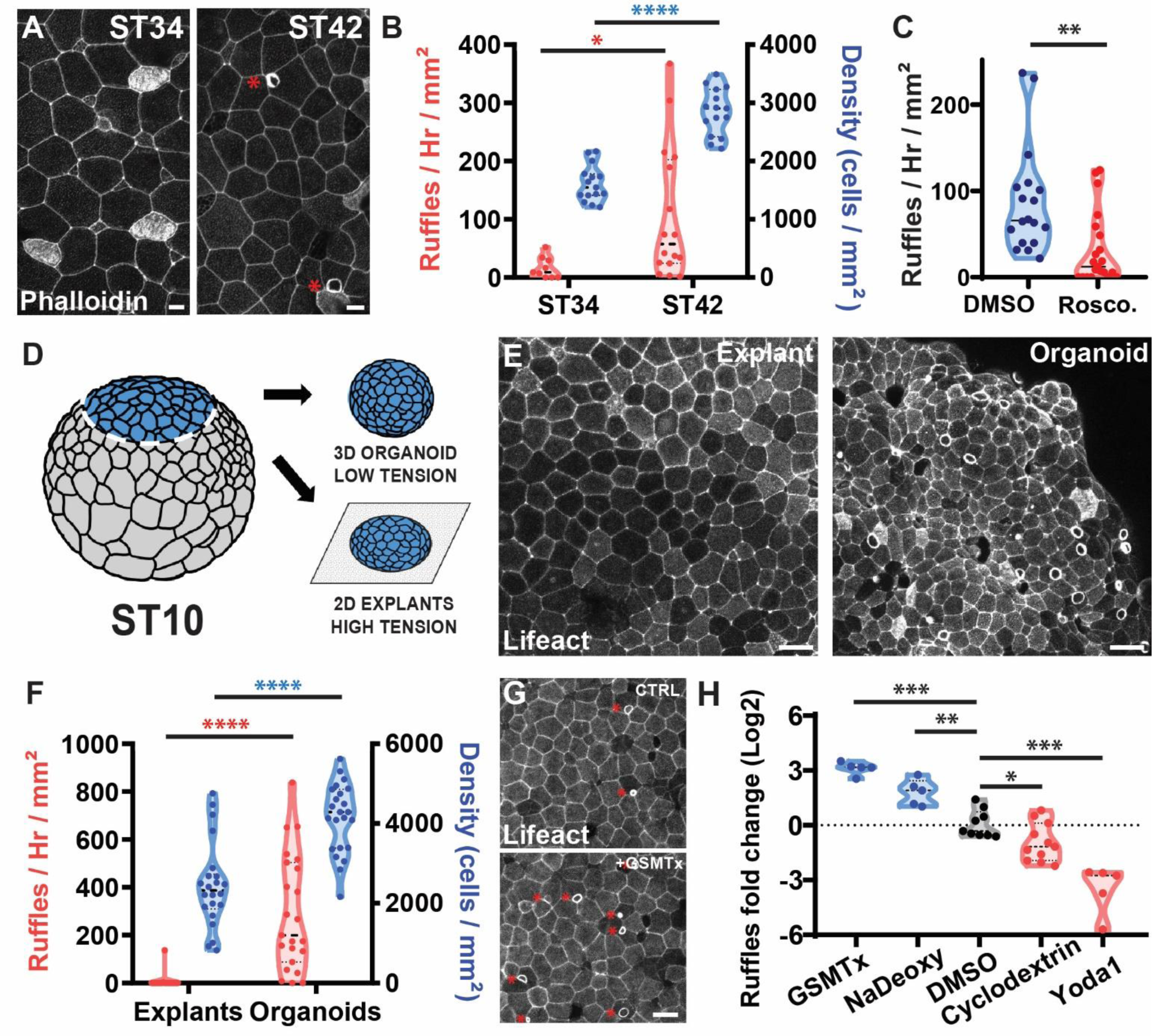
Macropinocytosis is associated with tissue crowding and low levels of tension. (**A**) Image of *Xenopus* stained with phalloidin (white) at ST34 (left) and ST42 (right, asterisk highlight actin ruffles). Scale bar 10 μm. (**B**) Quantification of cell density (blue; right axis) and macropinocytotic events (red; left axis) at ST34 and ST42 (n = 3 exp. and 15 embryos per stage for cell density, 4 exp. and 6 embryos for macropinocytotic events at ST34 and 5 exp. and 16 embryos at ST42). (**C**) Quantification of macropinocytotic events in ST44 embryos treated for 24H with either DMSO or Roscovitine (n = 4 exp. and 18 control embryos and 21 Roscovitine-treated embryos) (**D**) Representation of the generation of 3D organoids or 2D explant plated onto fibronectin coated coverslip. Adapted from ^47^. (**E**) Representative images of the surface of 2D explant (left) or 3D organoids (right) expressing Lifeact-FP. Scale bar 30 μm. (**F**). Quantification of cell density (blue; right axis) and macropinocytosis (red; left axis) in 2D explants or 3D organoids (n = 6 experiments with 22 explants and 23 organoids). (**G**) Image of actin ruffles (asterisk) in the epithelium of an embryo expressing Lifeact-FP before (up) and after (down) treatment with GSMTx4. Scale bar 30 μm. (**H**) Quantification of the fold difference in macropinocytosis level between 1Hr before and 1Hr after treatment with DMSO (mock treatment, n = 9 exp.), GSMTx4 (n = 5 exp.), NaDeoxy (n = 5 exp.), Mb Cyclodextrin (n = 11 exp.), or Yoda (n = 5 exp.).

To further investigate how tensile forces modulate macropinocytosis, we chemically manipulated membrane tension and assessed the consequence on macropinocytosis. First, we treated embryos with Mβ cyclodextrin, a small molecule known to remove cholesterol and lipid rafts from membranes that has been used to increase membrane tension ^28^, and we observed an ∼2-fold decrease in macropinocytosis **(Fig 2H)**. In contrast, we treated embryos with Sodium Deoxycholate (NaDeoxy), a detergent that at low doses can insert into the membrane and which has been used to lower membrane tension ^29^. In embryos treated with NaDeoxy we observed a significant ∼2-fold increase in macropinocytotic events consistent with our interpretation that low membrane tension promotes macropinocytosis **(Fig 2H)**.

Changes in membrane tension can be sensed via mechanosensory channels of the TRP and Piezo families. To confirm that mechanosensation is an important regulator of macropinocytosis in our system, we quantified macropinocytotic events in tissues where we manipulated mechanosensation both positively and negatively. Inhibition of mechanosensory channels (Trp and Piezo) via GsmTX4 ^30, 31^ led to an ∼3-fold increase in macropinocytosis **(Fig 2G, 2H)**. In contrast, activating Piezo1 with Yoda1 ^32^ led to a ∼4-fold decrease in macropinocytotic events **(Fig 2H)**. These results are consistent with findings in both human cells and the marine organism Hydra which have shown that macropinocytosis is negatively regulated by membrane tension ^14, 23, 33^. These results further suggest that tissue crowding induced macropinocytosis occurs via a decrease in tension.

### Macropinocytosis leads to cell size reduction

Various forms of endocytosis have been shown to participate in membrane remodeling during morphogenesis ^9, 11^. We therefore wanted to assess the consequence of macropinocytosis on the organization of the apical surface during epithelial development. By tracking the apical size of individual cells during macropinocytosis, we measured a loss of approximately 10% of apical surface area **(Fig 3A, 3B)**. While there is some variation to the change in cell size, we observe a correlation between the size of the macropinocytotic event and the reduction in cell size further supporting the idea that these events cause the reduction (r^2^=0.398, **Fig 3C**). Additionally, these events preferentially occur at cell-cell junctions, and we also observed a 10% decrease in the length of junctions associated with a macropinocytotic event **(Fig 3D)**. However, we did not observe the internalization of junction proteins as previously reported ^10^ **(Fig S1A-D)**. Therefore, we conclude that cell size reduction is primarily driven by internalization of a large portion of the apical membrane rather than internalization of the junctions *per se*.

**Fig. 3.**
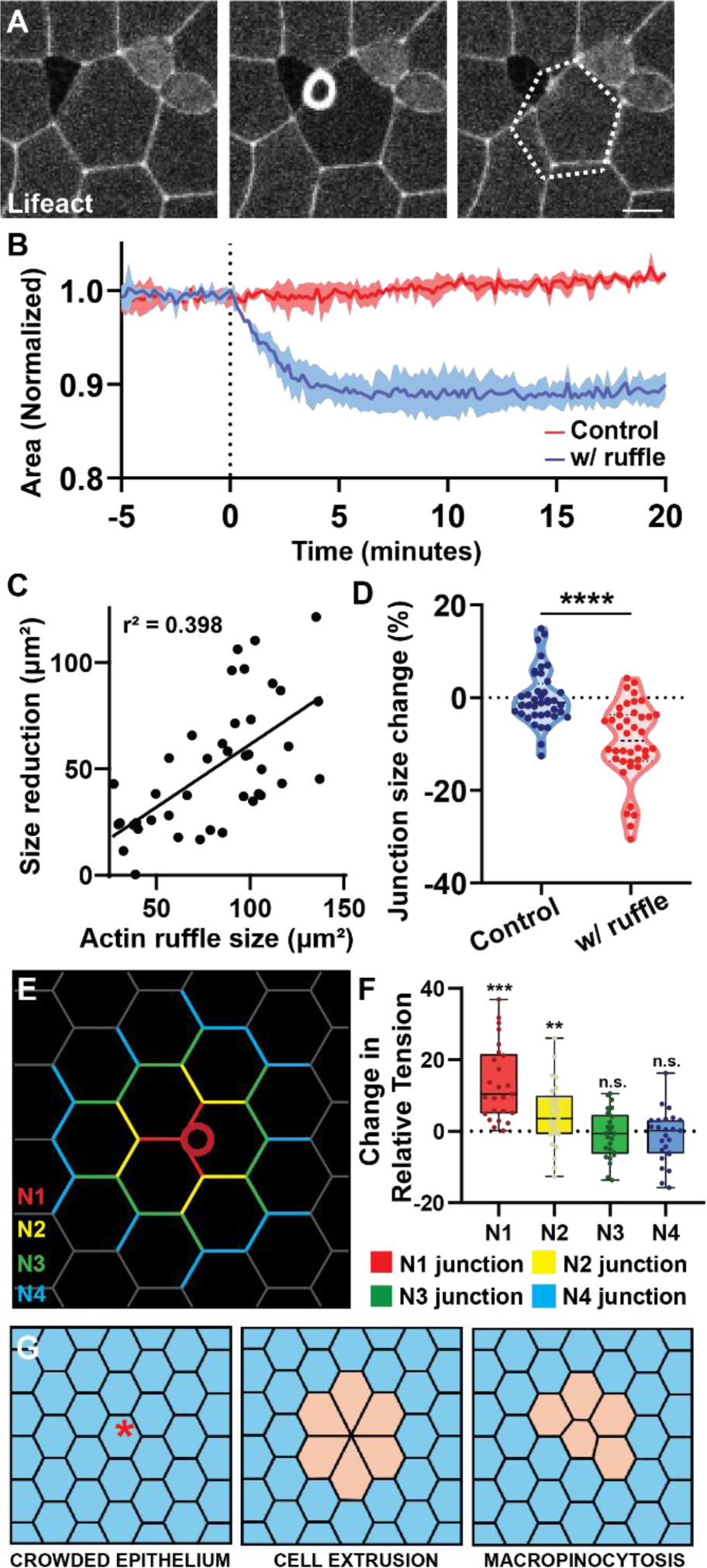
Macropinocytosis leads to cell size reduction. (**A**) Time lapse of a macropinocytotic event in tissue labeled with Lifeact-FP. The dotted line represents cell original size, highlighting the cell size reduction. Scale bar 10 μm. (**B**) Quantification of cell size change in cells with (blue) or without (red) a macropinocytotic event using cells synchronized to the onset of macropinocytosis and normalized to original cell size (n = 3 exp. and 43 cells). (**C**) Correlation between the apical size reduction and the size of actin ruffle (n = 3 exp. and 43 cells). (**D**) Quantification of the change in junction length in cells with and without macropinocytosis (n = 4 exp. and 40 cells). (**E**) Graphical depiction of a cell with different junction types labeled (N1-N4) in different colors based on their distance from the macropinocytotic event. (**F**) CellFit analysis of time lapse movies quantifying the change in relative junctional tension before and after macropinocytosis (n = 25 events; see Fig. S1). (**G**) Model comparing the effect of cell extrusion (center) or macropinocytosis (right) occurring in the cell (asterisk) of a crowded epithelium (left). Beige color highlights the putative effect on cells / neighbors.

Our observations of macropinocytosis-induced cell size reduction also revealed a local remodeling of the epithelium in response to the cell shrinkage, which should affect local tension distribution. To investigate this, we utilized the CellFit analysis that infers relative junctional tension based on tissue geometry ^34, 35^. We performed CellFit on frames before and after a macropinocytotic event **(Fig S1E)** and compared the changes to inferred tension in the junctions associated with a macropinocytotic event (N1), the junctions associated with the N1 junctions (N2) and so on **(Fig 3E)**. This analysis revealed that relative junction tension significantly increases after a macropinocytotic event for both the N1 and the N2 junctions, but that the effect dissipates by the N3 and N4 junctions **(Fig 3F)**. This local increase in junction tension suggests that cell size reduction by macropinocytosis induces a local stretching of the epithelium, similar, but weaker, than that observed during cell extrusion ^36^. From these results we propose that macropinocytosis can serve as a regulatory mechanism in response to crowding. Similar to cell removal by crowding-induced cell extrusion, cell size reduction by macropinocytosis would facilitate the alleviation of tissue crowding **(Fig 3G)**.

### Macropinocytosis regulates crowding during epithelial development

Live cell extrusion in response to tissue crowding has been observed in numerous tissues and organisms ^1, 2, 4^. We recently published that in *Xenopus* embryos, a subset of multiciliated cells (MCCs) are targeted for apical cell extrusion via mechanosensation. Importantly, this extrusion increases as the embryo develops suggesting that tissue crowding is a driving factor ^37^. Consistent with this, when we prevent crowding by inhibiting proliferation with Roscovitine, we observe a dramatic maintenance of MCCs compared to controls **(Fig 4A, 4B)**. Interestingly, while we have observed macropinocytotic events across the epithelium, we never observe them in MCCs, but instead have found a quantifiable bias for them to occur in cells adjacent to MCCs **(Fig 4C-E, Movie S1)**. We hypothesize that crowding-driven MCC extrusion could be affected by cell size reduction of neighboring cells via macropinocytosis.

**Fig. 4.**
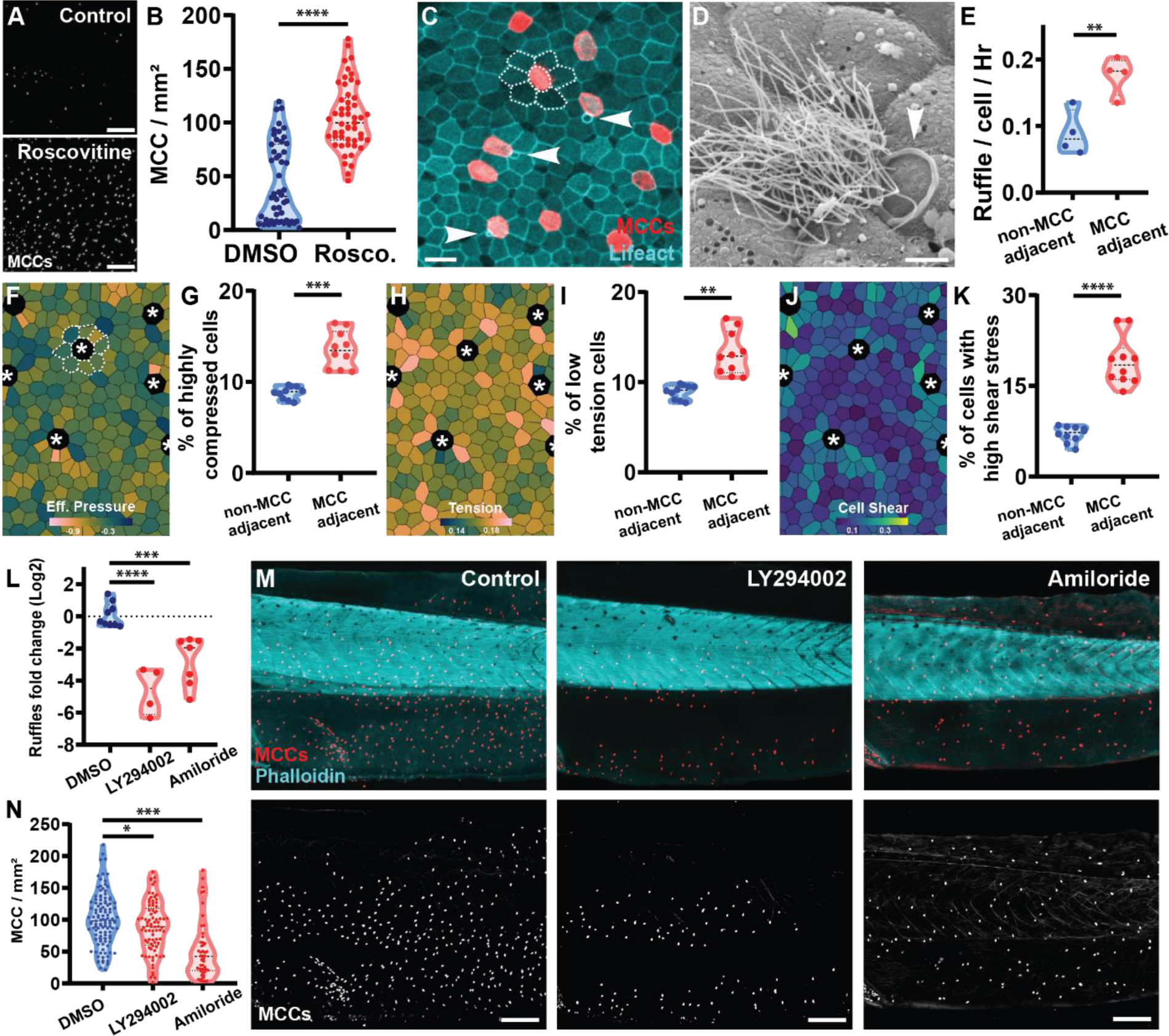
Macropinocytosis regulates tissue crowding during epithelial development. (**A**) Representative images of MCCs (marked with acetylated tubulin ab) in ST47 embryo after treatment with DMSO or Roscovitine. Scale bar 300 μm. (**B**) Quantification of MCC number in embryo treated for 24H with either DMSO or Roscovitine (n = 4 exp. and 50 DMSO and 51 Roscovitine embryos). (**C**) Representative image of macropinocytic events (arrowheads) occurring next to MCCs in the epithelium of a TubA1A-membrane RFP transgenic embryos labeling multiciliated cells (red) and injected with Lifeact-FP (cyan). Dashed lines represent rosette-like structure around MCCs. Scale bar 30 μm. (**D**) Representative SEM image showing a macropinocytotic event (arrowhead) adjacent to an MCC. Scale bar 5 μm. (**E**) Quantification of the number of events per cell per hour for cells adjacent or not adjacent to MCCs (n = 4 exp. 191 macropinocytotic events). (**F, H, J**) Images of a simulated epithelium with multiciliated cells (asterisk). Cells are color-coded for effective pressure (F), tension (H) or shear stress (J). Dashed lines represent rosette-like structure around MCCs. (**G, I, K**) Percentage of non-MCC neighbors and MCC neighbors that are in the lowest decile of effective pressure (G), tension (I) or shear stress (K), (n = 10 simulations). (**L**) Quantification of the fold difference in macropinocytotic events comparing 1Hr before and 1Hr after treatment with DMSO (n = 9 exp.), 1mM Amiloride (n = 7 exp.), or 5μM LY294002 (n = 4 exp.). (**M**) Representative images of MCCs (marked with acetylated tubulin ab; red / white) in ST46 embryos stained with phalloidin (cyan) after treatment with DMSO, LY294002 or Amiloride. Scale bar 300 μm. (**N**) Quantification of the number of MCCs after treatment with DMSO (n = 115 embryos), 1mM Amiloride (n = 81 embryos) or 5μM LY294002 (n = 49 embryos).

First, to understand why macropinocytosis is increased around MCCs we mathematically modeled the ciliated epithelium in order to explore how MCCs affect their neighboring cells. We defined MCCs as non-dividing cells with high stiffness (low deformability) due to the elaborate apical cytoskeleton needed to support the mechanical force associated with ciliary beating ^38-41^. Our simulations (**Movie S7, Fig 4F, 4H, 4J, Fig S2**) revealed that the presence of MCCs induce the organization of the surrounding cells into rosette-like structures (**Fig 4F, Movie S7**), resembling those observed *in vivo* (**Fig 4C**). Additionally, our model predicts that these rosettes are enriched in highly compressed cells (strongly negative effective pressure, **Fig 4F, 4G, Movie S7**) with lower tension levels (**Fig 4H, 4I, Movie S7**) compared to non-MCC neighboring cells. Similarly, MCC neighbors are enriched in cells with high shear stress (**Fig 4J, 4K, Movie S7**). These simulations suggest that high compression and low tension levels are responsible for macropinocytosis enrichment around MCCs, consistent with our experimental observations.

To properly investigate how local macropinocytosis around MCCs could affect the timing of their extrusion, we utilized two known small molecule inhibitors of macropinocytosis, Amiloride and the PI3K inhibitor LY294002 ^42, 43^. We found that, consistent with the literature, both drugs significantly reduced macropinocytosis in our system **(Fig 4L)**. Furthermore, extended treatment with either compound leads to premature extrusion of MCCs from the epithelium **(Fig 4M, 4N)** suggesting that, in the absence of macropinocytosis, tissue crowding is accelerated requiring a concomitant increase in cell extrusion. These results suggest that cell size reduction via macropinocytosis can alleviate tissue crowding which, as evidenced by the modulation of MCC extrusion, can have a profound impact on overall tissue architecture.

### A wave of macropinocytosis remodels the epithelium in response to external tissue compression

While the compressive stress of tissue crowding during development arises from proliferation, externally applied physical forces can also alter the compressive stress experienced by a tissue. In fact, epithelial compression has been reported to result in lower membrane tension ^3^. This led us to investigate if macropinocytosis could also provide a response to compression-induced stress. To investigate this, we first embedded embryos in agarose gel, similar to what has been used to induce compression in zebrafish embryos ^44^. In embryos compressed in 2% agarose, we find a rapidly induced wave of macropinocytosis across the epithelium with almost every cell having an event **(Fig S3A, S3B)**. However, in this rigidly confined space, the embryos do not survive for extended periods of time. We therefore developed a second approach for compression where we pressed embryos using a coverslip within a defined imaging chamber. This led to the flattening (and compression) of only the round parts of the embryo (e.g., the belly) which allowed for local imaging of just the compressed tissue without the lethality of whole embryo agarose compression. This compression induced a similar wave of macropinocytosis **(Fig 5A, 5B, Movie S8)** that could be imaged for extended periods of time.

**Fig. 5.**
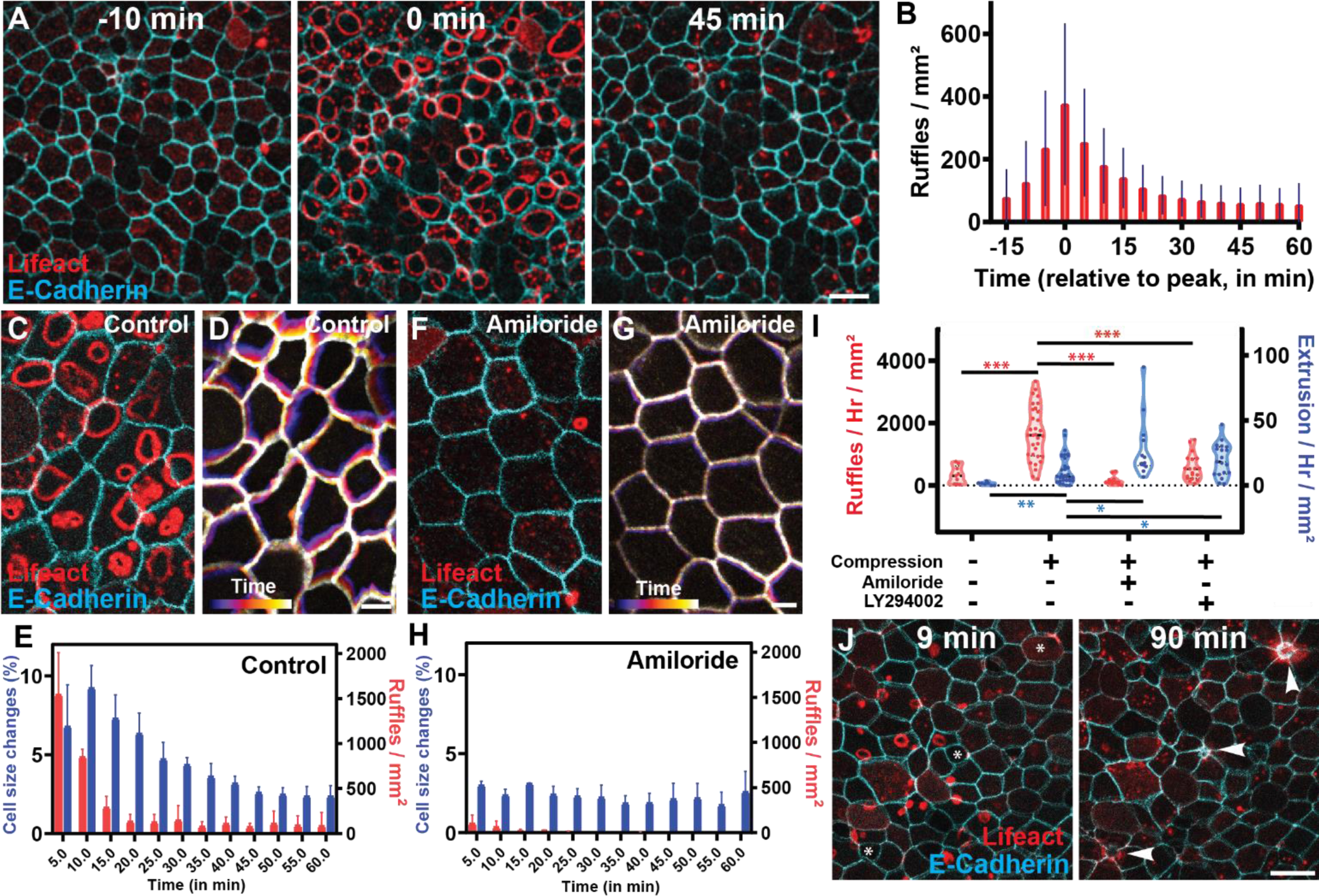
A wave of macropinocytosis remodels the epithelium in response to external tissue compression. (**A**) Representative image of an embryo injected with Cadherin-FP (cyan) and Lifeact-FP (red) before (left), during (middle), and after (right) a wave of macropinocytosis induced by tissue compression. Scale bar 30 μm. (**B**) Quantification of the wave of actin ruffles upon compression (n = 12 exp.). (**C, F**) Representative images of embryos injected with Cadherin-FP (cyan / white) and Lifeact-FP (red) 5 minutes after compression in control (C) or Amiloride treated (F) embryos. Scale bar 10 μm. (**D, G**) Representation of tissue remodeling over 10 min using a temporal color code of control (D) or Amiloride treated embryos. Scale bar 10 μm. (**E, H**) Quantification of tissue remodeling (i.e. overall change in cell size) (blue; left axis) and macropinocytotic events (red; right axis) in control (E) and Amiloride treated embryos upon compression (n = 3 exp.). (**I**) Quantification of macropinocytotic events (red; left axis) and cell extrusion events (blue; right axis) in non-compressed (n = 9 embryos) and compressed embryos in either control (n = 32 embryos) or embryos treated with amiloride (n = 13 embryos) or LY294002 (n = 19 embryos). (**J**) Representative image of an embryo injected with Cadherin-FP (cyan) and Lifeact-FP (red) having both macropinocytotic events and cell extrusion (arrowhead) upon compression. Scale bar 30 μm.

Because most cells contained a macropinocytotic event during this wave of macropinocytosis, we observed drastic tissue remodeling as a result of both cell reduction in the host cell and stretching in the neighbor cells. Quantification of cell size on isolated cells confirmed that macropinocytosis leads to a reduction in the apical surface area, similar to what we observed during epithelial growth **(Fig S3C)**. To evaluate overall tissue remodeling by the wave of macropinocytotic events, we measured absolute changes in cell size that considers both cell size reduction by macropinocytosis as well as cell stretching induced by events in the neighboring cells **(Fig 5C-E, Movie S8)**. We measured a peak in tissue remodeling that corresponds with the timing of the wave of macropinocytotic events **(Fig 5E)**. Additionally, in embryos in which we have inhibited macropinocytosis, we see a dramatic inhibition of epithelial remodeling via cell size change **(Figure 5F-H)**. These results indicate that macropinocytosis can serve as an important regulator of tissue remodeling not only during development but also in response to external stimuli.

As mentioned above, cell extrusion is an important response to tissue crowding and is known to occur in response to external compression ^1^. As expected, we observed a significant increase in cell extrusion events in response to embryo compression compared to uncompressed embryos **(Fig 5I, 5J, Movie S9)**. Importantly, embryos pretreated for 30 minutes with the macropinocytosis inhibitors exhibit a further increase in extrusion in response to compression **(Fig 5I)**. These results indicate that tissue remodeling associated with the wave of macropinocytotic events help to alleviate the effects of tissue compression. Importantly, in the inhibitor-treated embryos, an increase in cell extrusion compensates for the lack of macropinocytosis-induced cell size reduction. Overall, we propose that cell size reduction by macropinocytosis cooperates with cell removal by extrusion to counter developmental- or environmental-induced compression.

## DISCUSSION

We report a novel role for macropinocytosis as a regulator of compression in vertebrate epithelia. Maintaining homeostasis requires a constant balance between compression and stretching. An important response to compression is live cell extrusion, which locally stretches the apical area of the tissue. We propose that apical membrane internalization through numerous macropinocytotic events can serve a similar function of apical stretching by causing a cumulative loss of approximately 10% of the apical cell area from multiple cells, rather than the 100% loss of one cell during extrusion.

We observe this mechanism during development where proliferation leads to compression in the form of tissue crowding. We propose that a decrease in membrane tension induced by crowding triggers macropinocytotic events and that cell size reduction by macropinocytosis reshapes tissue architecture to alleviate crowding. The long-term importance of this mechanism for epithelial regulation is evidenced by the increase in cell extrusion when macropinocytosis is inhibited. Importantly, this mechanism is also utilized in acute response to external compression where a wave of macropinocytotic events extensively remodel the epithelium to alleviate the effects of compression. In this acute response, macropinocytosis and cell extrusion again cooperate to alleviate compression, with the inhibition of macropinocytosis being compensated for by an increased rate of cell extrusion.

This regulatory mechanism could also be at play in other vertebrates, including humans. Macropinocytosis and circular actin ruffling are widely conserved mechanisms reported in numerous human tissues. For example, similar actin-based internalization events have recently been reported in intestinal organoids ^45^. Additionally, macropinocytosis has been proposed to be regulated by tension in evolutionarily diverse organisms, from the marine organism Hydra to human cells in culture ^14, 23^. Finally, macropinocytosis and other endocytic structures have been shown to participate in both tissue remodeling and tension regulation ^9, 11, 12, 46^. Emerging evidence supports the broad potential use of macropinocytosis as a rapid and reversible response to compressive stresses.

Overall, this novel role for macropinocytosis offers an additional mechanism for vertebrate epithelia to regulate tissue crowding, particularly on relatively short timescales, and to continuously adjust tension distribution in response to stresses from both the environment and morphogenic growth. Our findings enhance the understanding of epithelial dynamics, highlighting how constant tissue remodeling is central to stress response.

## MATERIAL AND METHODS

### Transgenic Xenopus and mRNA embryo injections

*In vitro* fertilizations were performed using standard protocols ^1-4^ that have been approved by the Northwestern University Institutional Animal Care and Use Committee (IS00027128). Transgenic Xenopus expressing membrane-bound RFP driven by the tubulin promoter (Xla.Tg(tuba1a:MyrPalm-mRFP)^NXR^), were previously generated and obtained from the National Xenopus Resource Center (NXR) ^5^. Wild type or transgenic embryos were injected at the two- or four-cell stage with 50 pg mRNA. mRNA of Lifeact-GFP/RFP ^6^, ZO-1-RFP ^7^, E-Cadherin-GFP ^7^, SF9-GFP ^8^, membrane-RFP ^9^ were synthesized with the Sp6 mMessage Machine kit (Life Technologies, AM1340) and purified by RNeasy MiniElute Cleanup Kit (QIAGEN, 74204).

### Immunostaining

Embryos were fixed with 4% PFA/PBS. To visualize cilia, embryos were incubated with mouse anti acetylated α-tubulin (T7451; Sigma-Aldrich, 1:500) in 0.1%Triton/PBS for an hour at room temperature followed by Cy-2-conjugated goat anti-mouse secondary antibodies (Thermo Fisher Scientific) at 1:750 dilution in 0.1%Triton/PBS for two hours at room temperature. Phalloidin 568 (A12380, Invitrogen, 1:750) was used to visualize actin. Cell density was quantified manually using the FIJI software.

### Microscopy

Fluorescent light imaging was performed either using a Nikon A1R laser scanning confocal microscope (Figure 1A, 1B, 1D, 1E, 1F, 1H, 1I, 1J, 1K, 1L; Figure 2A, 2B, 2C, 2E, 2F, 2G, 2H; Figure 3A, 3B, 3C, 3D, 3F; Figure 4C, 4E, 4L; Figure 5A, 5B, 5C, 5D, 5E, 5F, 5G, 5H, 5I, 5J; Supplementary Figure S1A, S1B, S1C, S1D, S1E; Supplementary Figure S3A, S3B, S3C, S3D; and Supplementary Movies S1, S2, S3, S4, S5A/B, S7, S8) or a Nikon Ti2 microscope used as a widefield microscope (Figure 4A, 4B, 4M, 4N) or using a MizarTILT light sheet (Figure 1G; Figure 2H; Figure 4E, 4L) and a photometric Prime 95B camera. Objectives used included Plan Fluor 10x Ph1 DLL (NA 0.3), Plan Fluor 20x MImm DIC N2 (NA 0.75), Plan Fluor 40x Oil DIC H N2 (NA 1.3), Plan Apo VC 60x Oil DIC N2 (NA 1.4), and SR HP Apo TIRF 100xH (NA 1.49). Live embryos were imaged in anesthetic (0.01% Tricaine in 0.1X MMR).

Quantifications were done using FIJI Software ^10^. Cell density was quantified manually using the FIJI software. Cell size change tracking was performed using the FIJI TrackMate plugin coupled with Cellpose segmentation ^11^ on embryos co-expressing Cadherin-GFP for segmentation and Lifeact-RFP for ruffles visualization. Junction sizes were quantified manually using FIJI. To visualize dextran internalization, embryos expressing Lifeact-GFP were imaged in anesthetic with 0.125 μg/ml of 70 kDa TRITC-dextran (Chondrex). Quantification was performed using the linescan tool of FIJI. Quantification of adhesion protein internalization was performed on embryos expressing either Lifeact-RFP and E-Cadherin-GFP or Lifeact-GFP and ZO-1-RFP using the linescan tool of FIJI. Quantification of active myosin around the ruffles was performed on embryos expressing Lifeact-RFP and SF9-3xGFP using the linescan tool of FIJI. To quantify preferential macropinocytosis around multiciliated cell ruffles, ruffles were manually counted and classified based on whether they occurred in MCC-neighboring or non-MCC-neighboring goblet cells, and then divided by the total number of MCC-neighboring or non-MCC-neighboring goblet cells. Quantification of MCC density was performed on z-stacks in FIJI using the DoG detector of the TrackMate plugin from FIJI.

For SEM preparation, embryos were fixed for at least 24h in 2% glutaraldehyde on 0.1 M sodium cacodylate buffer at 4°C overnight and postfixed using 1% OsO4 in water for 2h. The specimens were then dehydrated in a graded series of ethanol (50%, 70%, 90%, and 100%), critical-point-dried with carbon dioxide (Samdri-790, Tousimis, Rockville, USA), mounted, and coated with 10 or 20 nm gold in a sputter coater. Finally, the specimens were observed under a scanning electron microscope (JCM-6000PLUS, JEOL, Japan).

### Drug treatments

To assess the effect of small molecules on the level of macropinocytosis, embryos expressing Lifeact-FP were first imaged for at least 30 minutes in anesthetic to establish their basal level of macropinocytosis. Subsequently, they were imaged for at least 45 minutes in anesthetic supplemented with either ^1^/_1000_ DMSO, 1 mM Amiloride (#14409, Cayman Chemical), 2 μM LY294002 (#HY-10108, MedChemExpress), 20 μM Blebbistatin (#B0560, Sigma), 0.25 mM Sodium Deoxycholate (#D6750, Sigma), 2 μM GSMTx (#HY-P1410, MedChemExpress), or 10 μM Yoda1 (#558610, Fisher). For cyclodextrin treatment, embryos were incubated for one hour in 50 mM Methyl-β-cyclodextrin (#J66847, Thermo) diluted in 0.1X MMR before being imaged for another hour in anesthetic. The level of macropinocytosis was manually quantified using FIJI, and results were expressed as log2(^treatment level^/_basal level_).

To assess the effect of macropinocytosis on MCCs extrusion, embryos were treated from stage 43 with either DMSO, 1 mM Amiloride, or 2 μM LY294002 in 0.1X MMR. Media were changed after 24 hours, and embryos were fixed in 4% PFA/PBS at 48 hours of treatment before processing for immunostaining. To assess the effect of proliferation on macropinocytosis or MCC extrusion, embryos were treated for 24 hours from stage 44/45 with either DMSO or 50 μM Roscovitine (#HY-30237, MedChemExpress) in 0.1X MMR. Subsequently, embryos were either imaged for 30 minutes to measure macropinocytosis level or fixed and processed for immunostaining to assess MCCs extrusion.

### Tension inference with CellFIT

To infer junction tension, embryos co-expressing Lifeact-RFP to visualize ruffle events and Cadherin-GFP for tissue segmentation were used. Tissue segmentation was performed using TissueAnalyzer with additional manual correction ^12^. The hand corrected segmented images were processed with Zazu CellFIT ^13^ to obtain infered tension. The results from 1-4 images before and after macropinocytosis events were averaged to obtain the final tension inference by junction category (N1, N2, N3, N4), expressed as a ratio (^average tension after macropinocytosis^/_average tension before macropinocytosis_).

### Compression assays

To assess the effect of agarose compression on macropinocytosis levels, embryos expressing Lifeact-FP were imaged in 2% agarose gel prepared in 0.5X anesthetic. For compression induced by coverslip, embryos under anesthesia were placed on a glass-bottom imaging dish (#P35G-1.5-14-C, Mattek), with a circle of grease applied outside the glass bottom, and an 18 mm coverslip (#72222-01, Electron Microscopy Science) placed on top of the grease circle. To examine the effect of macropinocytosis inhibition on compression-induced tissue remodeling, embryos expressing Lifeact-FP and Ecad-FP were pretreated with either ^1^/_1000_ DMSO or 1 mM Amiloride in 0.1X MMR for 30 minutes before imaging for 1 hour under coverslip compression. Cell size quantification was performed using the FIJI Trackmate plugin coupled with cellpose segmentation ^11^, while actin ruffles were manually quantified using FIJI software. To assess the impact of macropinocytosis inhibition on compression-induced cell extrusion, embryos expressing Lifeact-FP were pretreated with either ^1^/_1000_ DMSO, 1 mM Amiloride, or 2 μM LY294002 in 0.1X MMR for 30 minutes before imaging for 1 hour under coverslip compression. Manual quantification of macropinocytosis or cell extrusion was conducted using FIJI software. Effective internalization upon coverslip compression was assessed in embryos expressing Lifeact-GFP, imaged in anesthetic with 0.125 μg/ml of 70KDa TRITC-dextran.

### Ectodermal explants

To visualize macropinocytosis in explants and caps (organoids), animal caps from embryos expressing Lifeact-FP were dissected at stage 10 (ST10) and processed following methods described previously ^6^. Briefly, explants were either allowed to round up and cultured in 0.5X MMR or plated between a fibronectin-coated coverslip and an imaging glass-bottom dish (#P35G-1.5-14-C, Mattek), cultured in 0.5X DFA supplemented with BSA. Uncapped embryos served as a stage reference. Round-up and adherent explants were imaged at stage 34.

### Tissue simulations

Simulations were performed using the vertex model code developed in the authors’ previous work ^14^. This method uses incidence matrices to specify the topology of the vertex network ^15, 16^, allowing for direct manipulation and formulation of monolayer mechanics, and employing logarithmic relationships between forces and strains. Within the context of a vertex model simulation of an isolated epithelial monolayer, MCCs were modelled by focusing on their increased stiffness and lack of cell division. MCCs are distinguished from other cells by a prefactor, σ, multiplying the cell cortical tension and internal pressure. This means that a given difference between cell perimeter length and preferred perimeter length produces a tension force that is greater by a multiple equal to this prefactor for MCCs than it would be for bulk cells. Similarly, a given difference between cell area and preferred area gives a pressure force in MCCs greater than for bulk cells by a factor of σ. MCCs are also prevented from dividing, whereas bulk cells can divide when their age exceeds the cell cycle time. Cells were chosen to be MCCs by randomly selecting an initial cell within the monolayer, excluding cells at the monolayer periphery. This cell was made an MCC, and its neighbours, all of its neighbours’ neighbours, and all of their neighbours were removed from the pool of cells that could be chosen to be MCCs, ensuring that no path of fewer than 3 adjacent cells can be formed between any 2 MCCs. Another cell was then selected from the remaining pool of bulk cells, again excluding peripheral cells and those excluded by proximity to an existing MCC, and the process was repeated until there were no remaining bulk cells that had not been excluded.

For the data in this work, we produced 10 independent simulations for each parameter set. Monolayers were grown from an initial regular geometry of 61 cells to a size of 400 cells. At this point, a selection of cells was randomly chosen following the protocol described previously, and these cells were made MCCs. The stiffness of these MCCs was changed by a factor of σ, and they were no longer allowed to divide. The simulation was allowed to continue until the monolayer had grown to 800 cells, at which point all division was stopped and the monolayer was allowed to relax to equilibrium.

We ran simulations with σ=1, meaning MCCs are the same stiffness as bulk cells but do not divide, and σ=10. All of these simulations were repeated whilst allowing MCCs to divide, such that for σ=1, MCCs are entirely indistinguishable from bulk cells. For analyses, all cells at the periphery of the monolayer were excluded. All simulations were performed in the jammed regime with L_0_ = 0.75 and γ = 0.2, with uniform external pressure (0.5). Code and analysis scripts can be found in branch stiffer-cells3 of the GitHub repository ^17^.

## AUTHORS CONTRIBUTIONS

Conceptualization: EB, BJM

Methodology: EB, CR, OJ, BJM

Investigation: EB, EES, CR, OAH, AG, JS, JM, FK

Software: CR, OJ

Visualization: EB, CR, BJM

Funding acquisition: SW, OJ, BJM

Project administration: BJM

Supervision: CA, SW, OJ, BJM

Writing – original draft: EB, BJM

Writing – review & editing: EB, CR, EES, AG, JS, JM, SW, OJ, BJM

## COMPETING INTEREST DECLARATION

Authors declare that they have no competing interests.

**Sup. Fig. 1.**
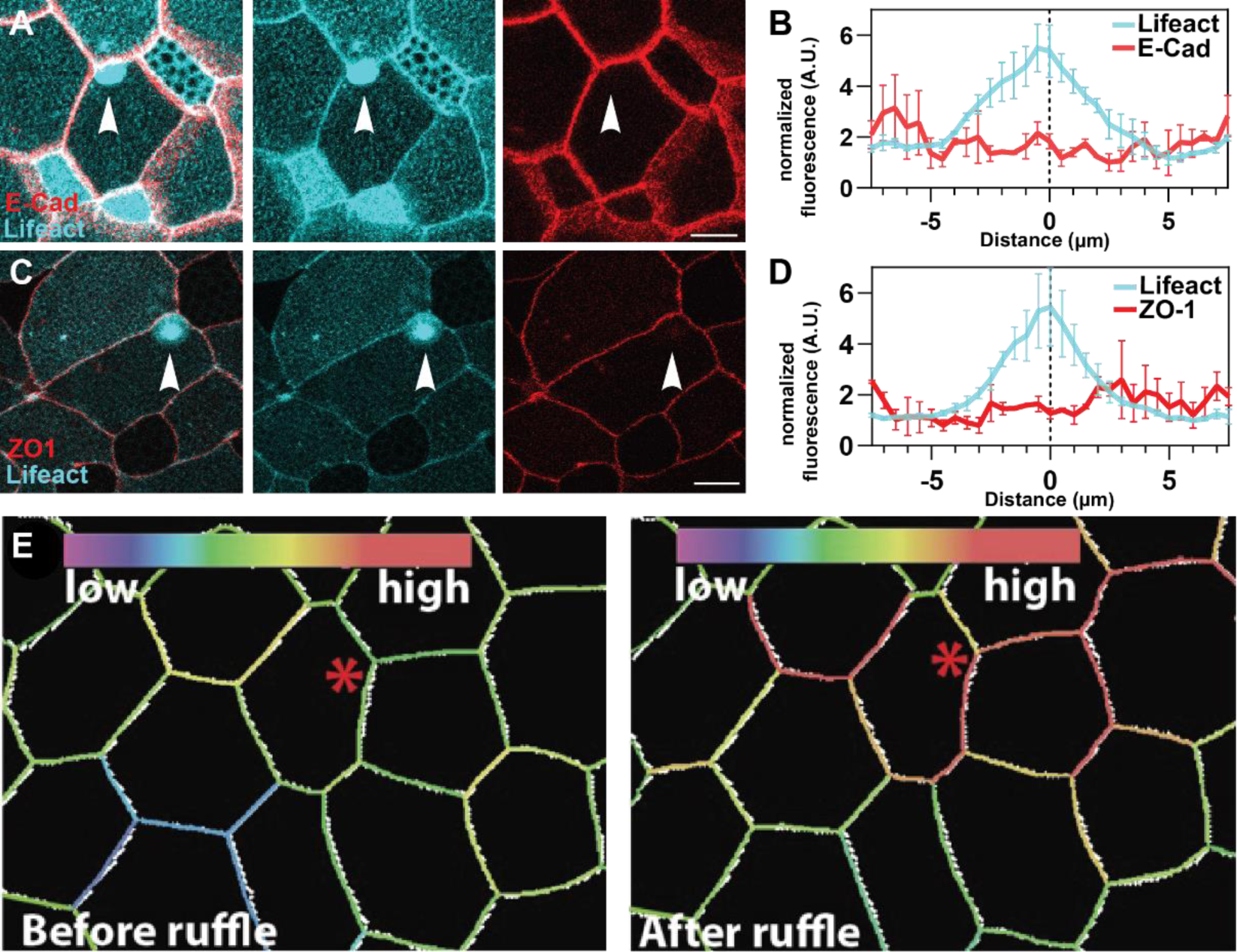
(**A, C**) Representative image of a tissue injected with (A) cadherin-FP (red) or (C) ZO1-FP (red) and Lifeact-FP (cyan) with a cell undergoing macropinocytosis. Scale bar 30 μm. (**B, D**) Quantification of the cadherin (B) or ZO1 (D) intensity associated with closing actin ruffles ((B) n = 3 exp. and 32 cells; (D) n = 2 exp., and 30 cells) (**E**) Inferred tension heatmaps using the CellFit program before (left) and after (right) a macropinocytotic event marked with a red asterisk.

**Sup. Fig. 2.**
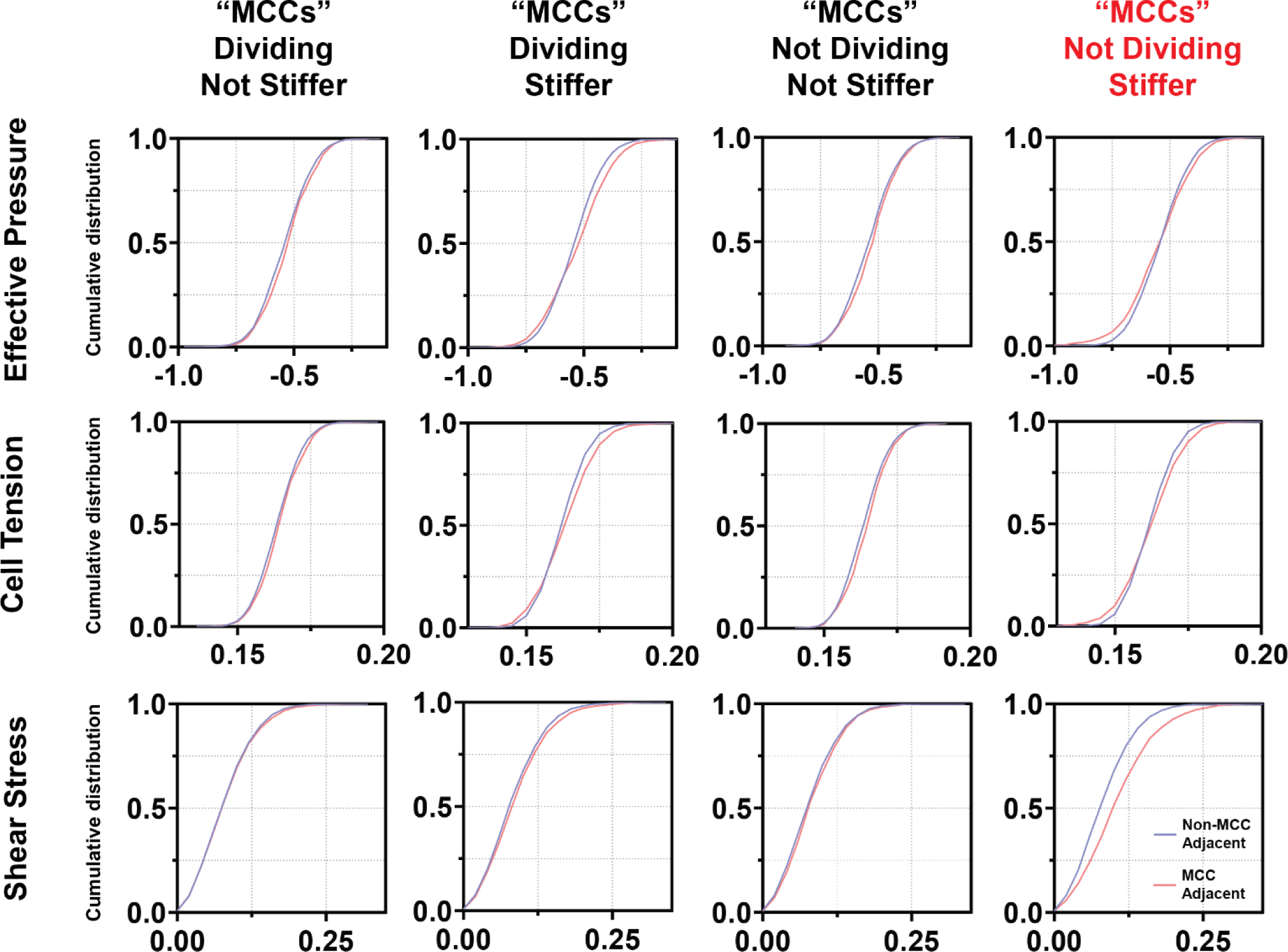
Cumulative distribution of cell’s effective pressure, tension or shear stress in simulated epithelium with multiciliated cells in name only (dividing and not stiffer), dividing and stiffer, not dividing and not stiffer, or not dividing and stiffer (n= 10 simulations per condition).

**Sup. Fig. 3.**
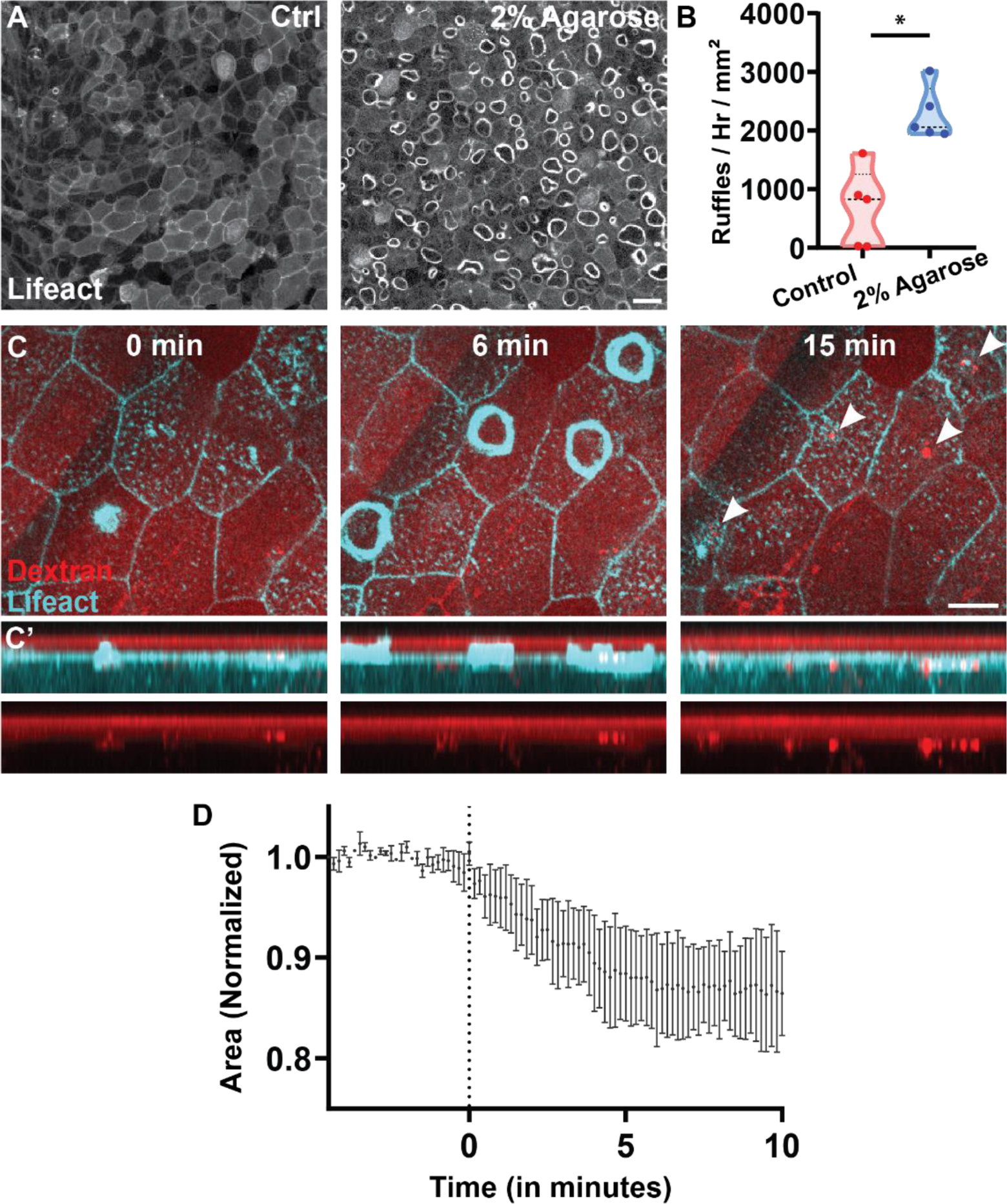
(**A**). Representative image of an epithelium expressing Lifeact-FP before (left) and after (right) compression in 2% agarose. Scale bar 30 μm. (**B**) Quantification of macropinocytosis before and after embryo compression in 2% Agarose (n = 2 exp. and 5 embryos). (**C**) Time lapse image of Lifeact-FP (cyan) showing macropinocytotic internalization of fluorescent dextran (red) from the media in embryo under compression. C’ show is a z projection of the area. Scale bar 10 μm. (**D**) Quantification of cell size change in tissues where we find isolated cells having a macropinocytotic event upon compression (n = 3 exp. and 45 cells).

